# Transcriptomics uncovers molecular mechanisms underlying salt-inhibited root growth in *Festuca rubra*

**DOI:** 10.1101/2025.07.22.665687

**Authors:** Bo Lan, Dafu Ru, Fang K Du, Kangquan Yin

## Abstract

Salt stress markedly inhibits root growth in *Festuca rubra*, while shoot development remains largely unaffected. Transcriptomic analysis identified 68,062 differentially expressed genes (DEGs) under salt stress. Upregulated genes were found significantly enriched in pathways related to methionine, melatonin, and suberin biosynthesis-processes likely contributing to stress adaptation through root growth suppression and maintenance of ion homeostasis. In contrast, genes associated with gibberellin biosynthesis, abscisic acid (ABA) response, and sugar signaling were downregulated, indicating complex hormonal and metabolic reprogramming. Notably, paradoxical regulation of gibberellin and ethylene signaling pathways suggests the presence of finely tuned mechanisms that balance growth and stress responses. These findings shed light on the molecular basis of root-specific salt stress responses in *F. rubra* and enhance our understanding of its adaptive strategies under saline conditions.

## Description

*Festuca rubra* L. (red fescue), is a versatile grass widely used as turfgrass and groundcover due to its fine texture and resilience. It is also effective for erosion control, as it stabilizes soil on slopes, along waterways, and in disturbed areas (Wright & Czapla, 2011). Additionally, red fescue has shown potential in phytoremediation, thanks to its ability to accumulate heavy metals such as copper, lead, manganese, and zinc from contaminated soils (Padmavathiamma & Li, 2009; Wong et al., 1994). Although occasionally used as pasture grass, it is not ideal for forage due to its limited nutritional value (USDA). Despite its high value and wide range of applications, red fescue also faces significant environmental challenges, including salinity. It is reported that approximately 20% of cultivated land and 33% of irrigated farmland worldwide are affected by high salinity, and without effective measures, over 50% of arable land may become salinized by 2050 (Jamill et al., 2011). Currently, most molecular insights into salt adaptation come from the model dicot species *Arabidopsis*, which is not inherently salt-tolerant, limiting the direct application of this knowledge to monocot red fescue (Diédhiou et al., 2009). Salt-stress response analysis in the halophytic red fescue *F. rubra* ssp. *litoralis* revealed that the expression of the *PIP2;1* aquaporin, V-ATPase subunit B, and the Na^+^/H^+^ antiporter *NHX1* was upregulated in roots under salt treatment. These genes were predominantly expressed in the root endodermis and vascular tissue, while expression was repressed in the epidermis and cortex under high salinity. This spatial regulation suggests that coordinated control of ion and water homeostasis contributes to salt tolerance. Further cDNA array analysis identified additional salt-responsive transcripts, including kinases and a WRKY transcription factor involved in signaling and gene regulation (Diédhiou et al., 2009). Nevertheless, the molecular mechanisms underlying salt-stress responses in red fescue remain largely unexplored.

In this study, we used the halotolerant red fescue variety *F. rubra* ssp. *rubra* ‘MAXIMA 1’ as a model to investigate salt stress responses. We treated the ‘MAXIMA 1’ with NaCl and compared the salt-stressed group with the control. The salt-stressed plants exhibited poor root development (Figure 1A), with significantly shorter root lengths compared to the control group (Figures 1A and B). In contrast, shoot length showed no significant difference between salt-stressed and control plants (Figure 1B). These results suggest that the roots of ‘MAXIMA 1’ are more sensitive to salt stress than the shoots, indicating a potential strategy in which halotolerant red fescue restricts root growth while maintaining shoot development under saline conditions.

**Figure 1.**
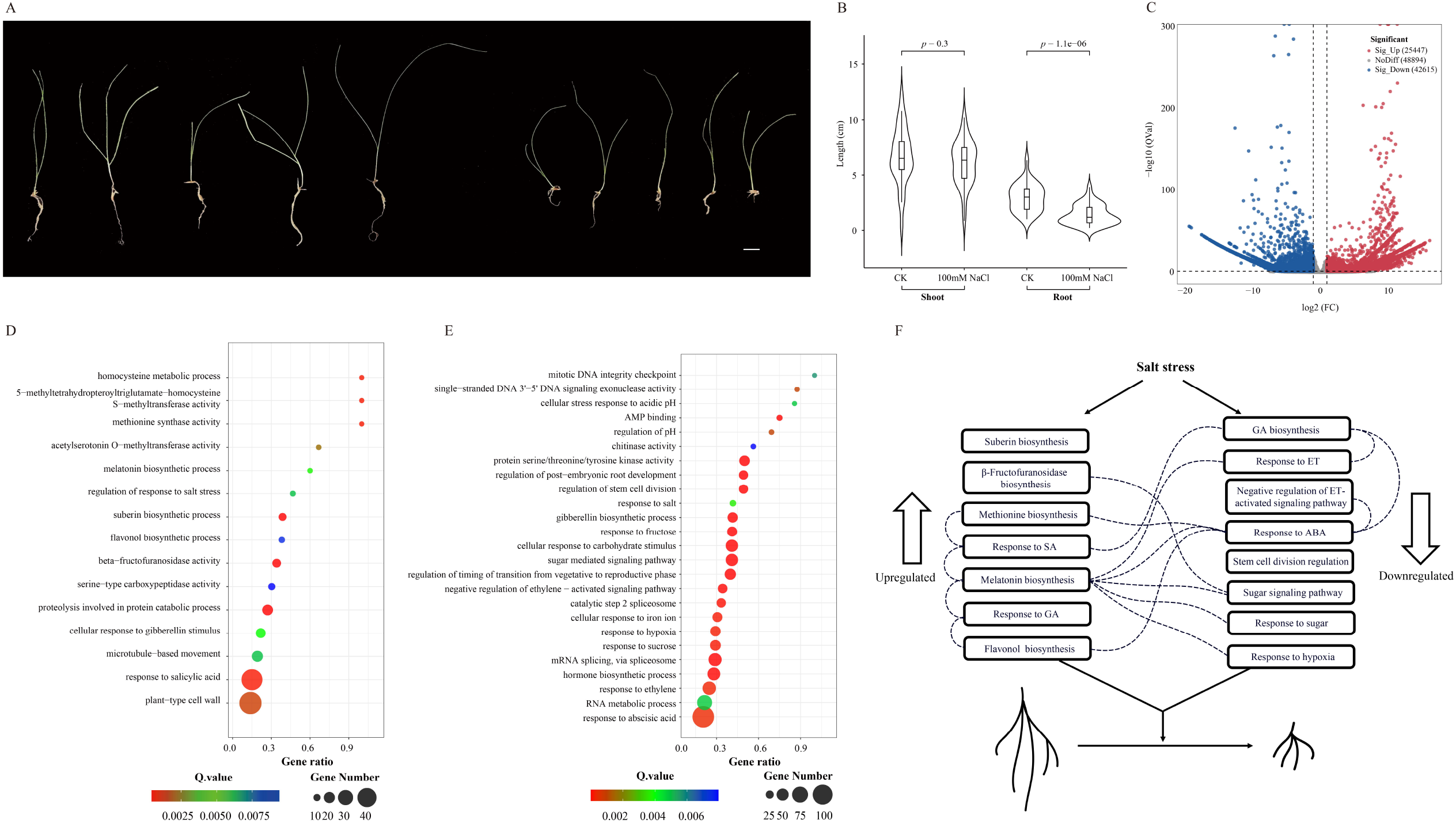
Phenotypic and Transcriptomic Responses of *F. rubra* to 100 mM NaCl Treatment. A. Growth performance of *F. rubra* on water-agar medium after 30 days. Left: control group; Right: plants grown under 100 mM NaCl treatment. Scale bar = 1 cm. B. Violin plots comparing shoot and root lengths between control and 100 mM NaCl-treated *F. rubra*. Independent samples t-test was performed for between-group comparisons, with p-values indicated in the figure. C. Volcano plot analysis of differentially expressed genes (DEGs). All unigenes from transcriptome data were included in the differential expression analysis. Each dot represents a unigene, with the x-axis showing log_2_ (fold change) (condition B/condition A) and the y-axis showing -log_10_(Q-value). Red dots indicate significant DEGs (|log_2_FC| ≥ 1 and Q-value < 0.05), while gray dots represent non-significant genes. Black dashed lines indicate thresholds for |log_2_FC| = 1 and Q-value = 0.05. D. Bubble plot of GO enrichment analysis for significantly upregulated genes (Q-value < 0.01). The x-axis shows the ratio of annotated gene number to background gene number for each term. Color scale represents Q-values, and bubble size corresponds to the number of annotated genes. E. Bubble plot of GO enrichment analysis for significantly downregulated genes (Q-value < 0.01). The x-axis shows the ratio of annotated gene number to background gene number for each term. Color scale represents Q-values, and bubble size corresponds to the number of annotated genes. F. Diagram illustrating *F. rubra*’s response to salt stress through transcriptional reprogramming. The left panel shows the functional categories of significantly upregulated genes, while the right panel displays those that are significantly downregulated. Dashed lines indicate potential interactions. Abbreviations: ABA, abscisic acid; GA, gibberellic acid; ET, ethylene; SA, salicylic acid.

To investigate the molecular mechanisms underlying the root-specific response of *F. rubra* to salt stress, we performed transcriptome profiling of root tissues from both control and salt-treated plants. Sequencing generated a total of 43.74 Gb of raw data, from which 285,284 transcripts were assembled, with a contig N50 of 899 bp and a GC content of 47.47%. From this assembly, 121,300 unigenes were predicted for subsequent differential expression analysis. A total of 68,062 genes were found to be significantly differentially expressed under salt stress, including 25,447 upregulated and 42,615 downregulated genes (Figure 1C). Among these, 39,760 genes were successfully annotated.

To investigate the functional implications of salt stress-responsive genes, we performed Gene Ontology (GO) enrichment analysis for differentially expressed genes (DEGs). Interestingly, among the upregulated genes under salt stress, GO categories related to methionine biosynthesis, melatonin biosynthesis, and suberin biosynthesis were significantly enriched (Figure 1D). Recent studies have shown that exogenous application of methionine can markedly enhance salt tolerance in several plant species (Shi et al., 2025). However, methionine has also been reported to inhibit primary root elongation in a dose-dependent manner, indicating a potential trade-off between growth and stress adaptation under high salinity conditions (Shi et al., 2025). This finding aligns well with our observations in *F. rubra*. Melatonin is a plant regulator known to enhance tolerance to various stresses, including salinity, by acting as a free radical scavenger to reduce membrane oxidation (Wang et al., 2018; Tahjib-Ul-Arif et al., 2025). As it promotes root growth at low concentrations but inhibits it at higher doses, we speculate that salt stress may induce elevated levels of melatonin in red fescue, thereby contributing to the inhibition of root growth (Figures 1A and B). The upregulation of suberin biosynthesis genes suggests increased suberin deposition in *F. rubra* under salt stress. Suberin accumulation in roots acts as a crucial barrier that helps reduce water loss and limits Na^+^ uptake (de Silva et al., 2021; Dabravolski & Isayenkov, 2023). This may help explain the differential growth between the above-ground and below-ground parts of *F. rubra*, as suberin can restrict Na^+^ accumulation in the roots, thereby reducing salt stress in the above-ground tissues. Moreover, we found that the GO category for beta-fructofuranosidase activity was significantly enriched among the upregulated genes in *F. rubra* under salt stress. This activity was also reported to be significantly upregulated in *Dunaliella salina*, an extremely halotolerant unicellular green alga, in response to hypersaline conditions (2.5M NaCl) (He et al., 2020), suggesting that both species may employ a similar strategy for salt tolerance despite having diverged over a billion years ago.

Among the downregulated genes under salt stress, GO categories related to gibberellin (GA) biosynthesis, negative regulation of the ethylene (ET) signaling pathway, and abscisic acid (ABA) responses were significantly enriched (Figure 1E). GA is a key hormone regulating cell proliferation and growth during plant development (Yamaguchi, 2008). In both dicots and monocots, GA-deficient mutants exhibit reduced root meristem size and shortened root length (Li et al., 2015; Griffiths et al., 2006), consistent with our observations in *F. rubra*. However, the GA response was upregulated (Figure 1D), presenting a paradox that suggests a compensatory mechanism maintaining appropriate hormone signaling to support growth and development. The downregulation of genes involved in the negative regulation of the ET signaling pathway may lead to pathway activation, which could further inhibit root elongation by affecting cell expansion (Růžička et al., 2007). Meanwhile, the ET response was downregulated—a paradox similar to that observed for GA, albeit in the opposite direction—suggesting a braking mechanism that limits overactivation of the ET signaling pathway (Figure 1E). Although salt stress generally activates the ABA signaling pathway (Geng et al., 2013), we found that ABA responses were suppressed in *F. rubra* under salt stress. This may represent a novel salt tolerance mechanism that could potentially be harnessed in other crops to generate resilient varieties. We also found enrichment of GO categories related to root development and stem cell division, which may result from GA deficiency or activation of the ET pathway. Interestingly, GO categories related to sugar signaling and response were enriched among the downregulated genes under salt stress. Although studies in model species have shown that salt stress generally induces sugar accumulation in plants, a comparative study of salt-sensitive and salt-tolerant rice varieties found that soluble sugar levels increased in the mature leaves of the salt-sensitive variety, but not in the salt-tolerant one (Pattanagul & Thitisaksakul, 2008). Based on these findings, we speculate that in *F. rubra*, sugars may be retained in the leaves to support shoot growth rather than being transported to the roots, thereby promoting above-ground development while suppressing root growth.

This study demonstrates that salt stress restricts root growth in *F. rubra* while maintaining shoot growth through transcriptional reprogramming. This involves the suppression of gibberellin biosynthesis, sugar signaling, and related responses, alongside the activation of ethylene signaling, amino acid, and melatonin biosynthesis. Structural changes such as increased suberization further enhance stress tolerance. Furthermore, multiple paradoxical patterns were observed, involving complex interactions between different hormones as well as feedback regulation within individual hormone pathways, underscoring the intricate and sometimes contradictory nature of hormonal signaling. These findings advance our understanding of *F. rubra*’s adaptation to salt stress and offer potential targets for improving stress resilience in crops.

## Method

### Salt Stress Treatment

*Festuca rubra* ssp. *rubra* ‘MAXIMA1’ seeds were purchased from Beijing Zhengdao Seed Co., Ltd. The seeds were soaked overnight and then surface-sterilized with 10% Clorox (sodium hypochlorite) containing 0.1% Tween-20 for 10–15 minutes, followed by thorough rinsing with sterile water. After drying, seeds for the control group (CK) were plated on water-agar medium, while seeds for the experimental group were plated on water-agar medium supplemented with 100 mM NaCl. Each treatment included three biological replicates, with 100 seeds per replicate. The seeds were incubated at 25 °C under a 12-h light/12-h dark photoperiod for 30 days. Root growth was monitored and photographed (Figure 1A) for further analysis. Root and shoot lengths were measured using ImageJ (Collins, 2007). Statistical significance between treatments was evaluated using Student’s t-test.

### Transcriptome Sequencing and Differential Gene Expression Analysis

RNA extraction from roots was performed using NEBNext Ultra RNA Library Prep kit (NEB, USA, Catalog #: E7530L). Paired-end RNA sequencing was performed on the Illumina NovaSeq6000 platform. We processed raw reads with Trimmomatic (Bolger, et al., 2014) to remove low-quality sequences and adapters. We assembled the transcriptome de novo using Trinity v2.6.5 (Haas et al., 2013) with default parameters, then evaluated assembly quality with contig N50 for continuity. We predicted protein-coding sequences using TransDecoder v5.7.1 (Hass, 2023) and we retained only the longest transcript per gene based on coding sequence (CDS) length manually. We performed functional annotation using EggNOG 5.0 (Huerta-Cepas et al., 2018) with the Gene Ontology (GO) database (Ashburner et al., 2000). To improve taxonomic relevance, the Taxonomic Scope parameter was set to Viridiplantae (Taxon ID: 33090). To avoid redundancy among GO terms, we employed a GOATOOLS-based (Klopfenstein et al, 2018) approach that utilizes the directed acyclic graph (DAG) structure of the Gene Ontology to remove redundant terms, retaining specific terms that are not ancestors of any other terms in the set. We quantified aligned reads using featureCounts v2.0.3 (Liao et al., 2014) and normalized expression levels to Transcripts Per Million. For differential expression analysis, we applied the DESeq2 pipeline (Love et al., 2014), defining significantly DEGs as those with |log_2_FoldChange| ≥ 1 and padj < 0.05 (Figure 1C), controlling the false discovery rate via the Benjamini-Hochberg method (Benjamini & Hochberg, 1995). We used all functionally annotated genes as the background set for GO enrichment. GO enrichment analysis was performed on the OmicStudio platform (Lyu et al., 2023), with terms considered significantly enriched at padj < 0.01.

## Data availability

We have deposited the transcriptome raw sequencing data in the NCBI Sequence Read Archive (SRA) under the BioProject accession number PRJNA1264723.

## Acknowledgements

We thank Yuqing Jing for collecting samples for RNA-seq.

## Author Contributions

KY conceived the study and designed the experiments; BL performed the experiments. BL, KY, FD and DR analyzed the data; BL and KY wrote the manuscript.

